# phylostratr: A framework for phylostratigraphy

**DOI:** 10.1101/360164

**Authors:** Zebulun Arendsee, Jing Li, Urminder Singh, Arun Seetharam, Karin Dorman, Eve Syrkin Wurtele

## Abstract

**Motivation:** The goal of phylostratigraphy is to infer the evolutionary origin of each gene in an organism. Currently, there are no general pipelines for this task. We present an R package, `phylostratr`, to fill this gap, making high-quality phylostratigraphic analysis accessible to non-specialists.

**Results:** Phylostratigraphic analysis entails searching for homologs within increasingly broad clades. The highest clade that contains all homologs of a gene is that gene’s phylostratum. We have created a general R-based framework, `phylostratr`, for estimating the phylostratum of every gene in a species. The program can fully automate an analysis: select species for a balanced representation of each strata, retrieve the sequences from UniProt, build BLAST databases, run BLAST, infer homologs for each gene against each subject species, determine phylostrata, and return summaries and diagnostics. `phylostratr` allows extensive customization. A user may: modify the automatically-generated clade tree or use their own tree; provide custom sequences in place of those automatically retrieved from UniProt; replace BLAST with an alternative algorithm; or tailor the method and sensitivity of the homology inference classifier. `phylostratr` also offers proteome quality assessments, false-positive diagnostics, and checks for missing organelle genomes. We show the utility of `phylostratr` through case studies in *Arabidopsis thaliana* and *Saccharomyces cerevisiae*.

**Availability:** `phylostratr` source code and vignettes are available on GitHub at https://github.com/arendsee/phylostratr

## 1 Introduction

The diversity of species on earth rises as species adapt to changing environments through the continual modification and re-purposing of existing genes and the emergence of new genes. Phylostratigraphy is the process of determining the phylogenetic origin of every gene in a genome [1]. Phylostratigraphy classifies genes by age, where age is defined relative to the branch on the tree where the gene is inferred to have first appeared in a genic state.

Phylostratigraphy has been applied to infer the novel proteins recruited in the evolutionary emergence of new structures and functions. For example, the over-represented expression of genes from the chordate phylostratum in the midbrain has been used to infer the genes involved in the origin of this structure [2]. Also the unique expression of an orphan (species-specific) gene in the shoot apical meristem of *Arabidopsis* has led to conjecture on its role in this structure [3]. By linking genes to their point of origin, we can identify candidate genes for involvement in clade-specific pathways [4].

Phylostratigraphy is also a first step towards understanding the evolutionary paths a gene may follow as it evolves from non-genic sequence or as a completely novel reading frame within an existing gene [5]. The changes that a young gene has undergone can be inferred by analysis of all the members of a lineage-restricted family. For example, the Antarctic notothenioid fish gained an anti-freeze protein, not observed in any other species, through the tandem duplication of a glycotripeptide [6]. Speciation is thought to occur in part by emergence of new genes with novel functions that permit organisms to respond to changing environments [4]. The early stages in evolution of a new protein-coding gene can be revealed by comparing the sequences of that gene across distinct populations of the species. Very new genes, termed “orphan genes”, encode species-specific proteins with no identifiable homologs in other species.

While phylostratigraphy is conceptually simple, it is in practice difficult for several reasons. Biological nuances – such as domain shuffling and horizontal gene transfer – can complicate inferences. Further, phylostratigraphy is highly sensitive to false positives among the inferred homologs, and tools for homology inference are far from perfect. Sequence data may be incomplete or inaccurate, gene annotation may be poor, and phylogenetic trees may be incorrect or ambiguous (i.e., there may be many unresolved nodes). The data can be systematically biased: some phylostrata may be over-represented, organelle-encoded proteins may be unavailable, and approaches to gene annotation may differ across time and by species. Finally, there are the mundane, time-consuming complexities of implementing the pipeline, such as acquiring sequence data, building the databases, and managing the proteome files and their phylogenetic relationships. All these factors can introduce errors in phylostratigraphic designations that cannot be easily caught. These challenges highlight the importance of diagnostics in phylostratigraphic analysis.

There are currently no phylostratigraphy pipelines that offer a high-degree of automation along with customization and powerful diagnostics. There is a web-based tool for phylostratigraphy, ORFanFinder [7]. However, analysis using ORFanFinder is limited to the strata corresponding to the named ranks of the NCBI common tree, which is too low-resolution for many purposes. Also, ORFanFinder does not provide flexible diagnostics or analysis beyond an immediate inference of gene ages. A few other published scripts are available for phylostratigraphy, but these are tailored to the needs of the particular study (e.g., [8]).

Here, we describe `phylostratr`, a customizable R package for reproducible, whole-genome phylostratigraphy. `phylostratr` automates what can be automated (e.g., sequence retrieval, database building, file handling, and NCBI tree building) and makes customizable what may need to be customized (e.g., input clade trees, homology classifiers, and protein sequence data). The package provides a suite of tools for diagnosing common pitfalls, catching phylostratigraphic oddities, and preparing output data for specialized analysis. We envision `phylostratr` being used as a flexible pipeline for phylostratigraphic annotation of genomes and as a framework for downstream analysis of the emergence and development of genes.

## 2 Implementation

`phylostratr` is an R package [10] distributed under the GNU public license. It is dependent on the NCBI BLAST+ standalone program [11], the R packages `taxizedb` [12] for NCBI taxonomy access, `ape` for tree representation, `dplyr` for table manipulations [13], and `ggplot2` and `ggtree` for visualization [14,15].

## 3 Approach

The purpose of `phylostratr` is to make phylostratigraphic analysis accessible, introspective, reproducible, and flexible. We make the pipeline accessible by automating much of the complex data handling (e.g., tree operations and proteome sequence retrieval) and sequence database tracking. `phylostratr` is introspective in that it provides powerful tools for finding flaws and irregularities in both inputs and results. `phylostratr` pipelines are reproducible since the entire pipeline, from data retrieval to diagnostics to downstream analysis, can be written and shared as a single R script. Flexibility is achieved by implementing the tools as a library within the powerful R analytic environment allowing the seven core functions in `phylostratr` to be easily customized. In the following sections we walk through the core phylostratigraphic pipeline (**Figure 1**) and discuss our reasoning behind unconventional choices.

**Figure 1.**
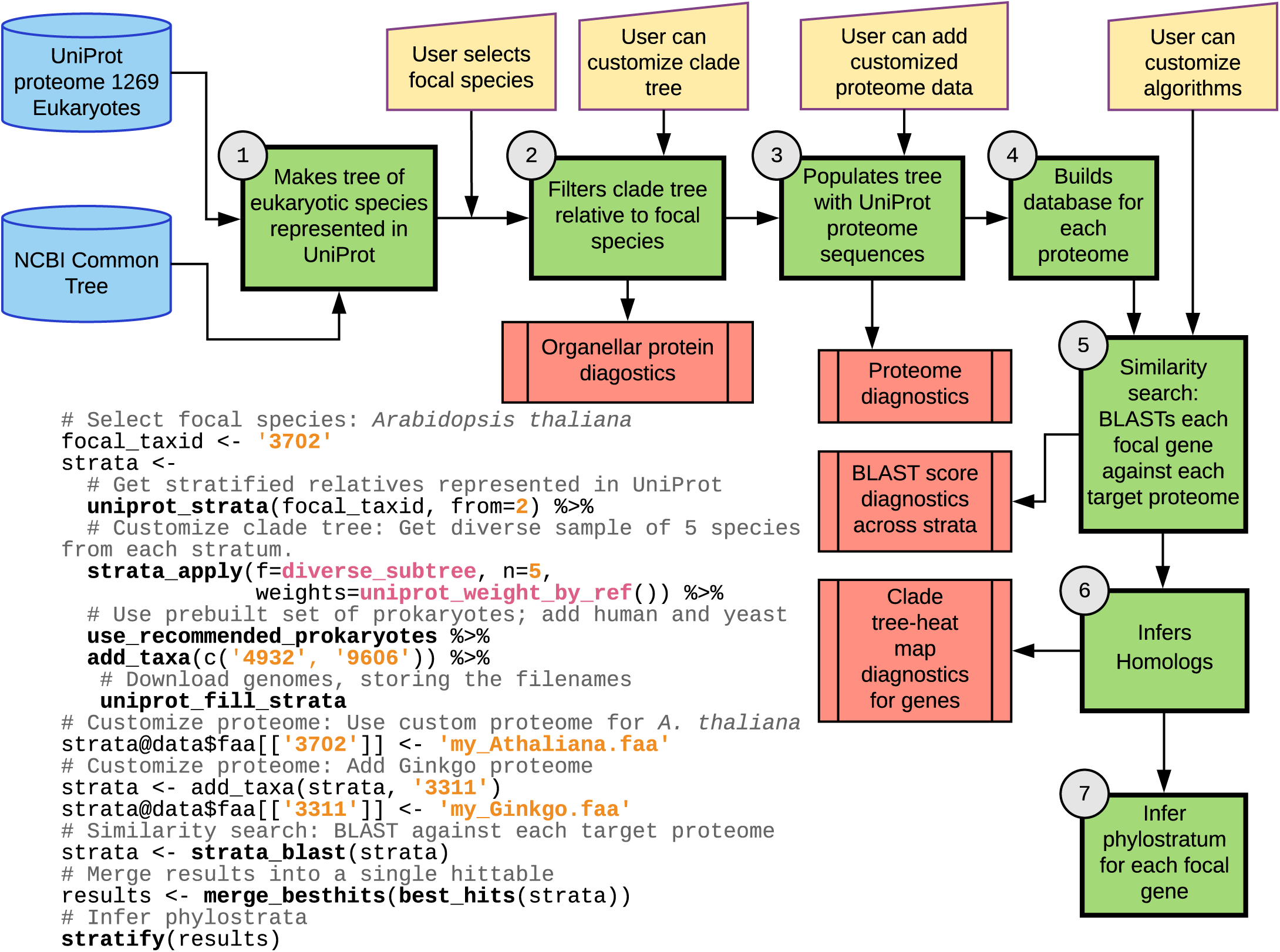
Overview of `phylostratr`. **Flow chart:** 1) `phylostratr` creates a clade tree from the species currently represented in UniProt, based on the current NCBI tree of life. 2) It then makes a trimmed clade tree that is centered around the focal species, using an algorithm designed to maximize the evolutionary diversity of species while honoring the constraints given by the user. 3) This diversified, trimmed clade-tree is populated by the proteomes from UniProt Proteome [9]. `phylostratr` automatically tracks the hundreds of species and thousands of annotated proteins/species that are used in each analysis. 4) It builds a database for each proteome and 5) performs a pairwise BLAST of the focal species proteome against the proteome of each species in the clade-tree (for greater speed, this search can be run on a distributed server). 6) The “best hits” for each focal gene against each target species are found and passed to a function that 7) assigns each gene to the phylostratum associated with the deepest clade to which the gene has an inferred homolog. This entire pipeline is run automatically by `phylostratr`. The focal species (3702) is the only input required for the entire analysis. Alternately, the user can customize the analysis. Innovative diagnostics are available. Green boxes, R core phylostratigraphy functions. Yellow trapezoids, customizable features. Salmon boxes, final output and diagnostics. **R code, customized to execute pipeline for *A. thaliana*:** The maximum number of species to be included in each clade are specified (n=5), this number may be lower if there are not five representatives. The user may add weights to the species-selection algorithm, here we set a preference for UniProt reference species; two eukaryotic species were added to the tree (NCBI taxon IDs 4932 and 9606), and a set of 85 user-selected prokaryotic species were added. The default UniProt proteome for the focal species was replaced with a custom one ‘my_thaliana.faa’. An initial analysis of *A. thaliana* in `phylostratr` showed that *Picea alba*, the only representative of the Spermataphyta phyloclade in Uniprot, had a sparsely annotated genome; thus, the Ginkgo proteome from the 1KP project was added.

### 3.1 Create a full clade tree

The first core function in `phylostratr` creates a large clade tree by mapping all species in the UniProt proteomes database [9] to the NCBI common tree [16]. The UniProt Proteome database includes only species with sequenced genomes.

### 3.2 Build trimmed clade tree with high-quality representatives for each phylostratum

The next core function prunes the giant clade tree such that it retains only select representatives for each phylostratum. To achieve this, `phylostratr` provides a novel algorithm that selects a diverse set of species from a tree while respecting a weight vector given by the user. The user provides an initial weight for each species (by default all weights are 1). `phylostratr` then loops through the following two steps until either all species in a phylostratum have been selected or the threshold number of species is reached. Step 1: the species with the highest weight is selected, with ties resolved by depth in the tree (preferring the deeper node). Step 2: the weights of all species are adjusted to penalize relatedness to other species. The number of previously selected species descending from each node in the tree is counted. A diversity score for each species is calculated by summing these counts from root to leaf. The diversity-penalized weight is the initial weight divided by the diversity score. Overall, in the unweighted case, this algorithm will always select the species that has the fewest internal branches in common with other selected species.

This selection method selects one deeply nested member from each stratum, but then prefers the more deeply rooted taxa in a stratum. Assuming an ultrametric phylogenetic tree, where all leaf nodes are the same evolutionary distance from the root node, more of the total evolutionary time traversed by the taxa in the stratum will be covered by selected descendent taxa and few very closely related species will be selected. In short, the selected taxa will be a diverse representation of the stratum. This preference for evolutionary diversity can be extended to non-ultrametric trees, by computing branch length-weighted counts of selected species in step 2 of the algorithm.

The user may modify this sampling strategy at each step. The NCBI common tree may be replaced (all or in part) with a custom tree; this is particularly important in the resolution of polytomies (unresolved lineages with multiple branches from a single node), which are common in the NCBI tree (**Figure S2-4**). Also, the species sampling step can be modified by altering the weights in the sampling algorithm, for example, to favor inclusion or exclusion of particular species or to set a preference for the UniProt reference proteomes (as shown in the R code of **Figure 1)**. The species sampling step can be omitted entirely, in which case all proteomes in UniProt would be included in the analysis.

Our choice to limit the search space to a discrete number of proteomes is somewhat unusual. It is common practice in phylostratigraphy to search for homologs in the largest database possible (usually the NCBI `nr` database). We deviate from this convention for two reasons. First, *using a discrete set of proteomes allows clear inferences not only about where a gene has a homolog*, *but also about where it does not have one.* This provides insight into the evolutionary history of each gene. Second, *limiting the search space reduces the chance of observing a false positive.* Any phylostratigraphic classification depends solely on the single most distant inferred homolog, thus it is sensitive to false positives. These false positives may arise by chance or by convergent evolution (either by selection for functional motifs or neutral expansion of repeats). As the number of representative species for a stratum grows, the false positive chance rises while the chance that the true homolog is either missing or altered beyond recognition decreases.

### 3.3 Populate clade tree with proteome

`phylostratr` will automatically download proteomes for species that are represented in the UniProt database (third core function). The user can input their own sequence data, in addition to, or in place of, the UniProt data. `phylostratr` offers a suite of tools for diagnosing gene histories and common problems with input proteomes (discussed in **Section 4)**.

### 3.4 Build proteome databases

For handling subsequent similarity comparisons, `phylostratr` creates databases from the proteome sequences. There are three possible ways to partition the sequences into databases. 1) all the sequences could be placed in a single pooled database; 2) sequences can be partitioned into one database per phylostratum; or, 3) each species could have its own database. The partitioning approach determines how the E-value (the expected number of hits with greater or equal score) is interpreted. The first way is by far most common in literature. However, we chose the third option – each species having its own database – since it allows us to interpret E-values for each species without a statistical dependence on other species in the analysis. The approach allows us to use sequence similarity tools other than BLAST, such as HMMER [17]. Also important is that since each species has its own database, species may be added or removed from the study without affecting previously computed E-values. This enables the species selection to expand as proteome sequence for additional species becomes available, without having to redo the whole analysis. This functionality is key to scalability.

### 3.5 Search for similarity

The next core function in `phylostratr` entails a similarity search between each focal genes and all proteins from all species in the proteome database. BLAST is the most popular similarity search algorithm, and is the default tool for phylostratigraphy [18]. `phylostratr` offers dedicated handling for BLAST: it automatically builds BLAST databases for all proteomes, runs BLAST alignment, and merges the results. Alternatively, the user may opt to run the BLAST search on a distributed system and then feed the results back into `phylostratr`.

There is some debate about the efficacy of BLAST to infer distant gene relatives, especially when functional constraints (and thus site-specific mutation rates) are not constant [18–21]. The `phylostratr` user may replace BLAST with a different algorithm for finding sequence similarity, for example to determine similarity across distant genes. One possibility is HMMER, a profile Hidden Markov Model (profile HMM) algorithm for sequence homology inference [17]. HMMER has been used to supplement phylostratigraphy studies of particular gene families (e.g [22]), though it has been applied only rarely for whole-genome phylostratigraphy (e.g. [23]). To change the similarity algorithm, the user writes a new classifier function and replaces the BLAST wrapper with a wrapper for the replacement algorithm.

### 3.6 Infer homologs based on similarity

Homology is inferred from the similarity results. In most phylostratigraphic studies this entails selecting an E-value homology threshold for a BLAST search of query protein against a pooled database of proteins from many species. All hits to the query protein with E-values below the cutoff are recorded and the significant hit in the most phylogenetically distant species determines the phylostratum classification.

We frame the phylostratigraphy challenge in a hypothesis framework that more closely fits the structure of the problem. Specifically, we want to test the hypothesis that a query gene has a homolog in a given phylostratum. In order to achieve this, we convert each E-value of the query gene against each target species to a p-value, adjust the p-values for multiple testing, and then return the most significant hit in each phylostratum. The default p-value adjustment method is the Holm method, though we use the more stringent Bonferroni method in our case studies. The reason we use Bonferroni in our case studies, is to more easily compare to past results based on pooled BLAST databases, which implicitly perform a Bonferroni correction. We compare results from Bonferroni versus Holm method in (**Figure 2A**).

**Figure 2.**
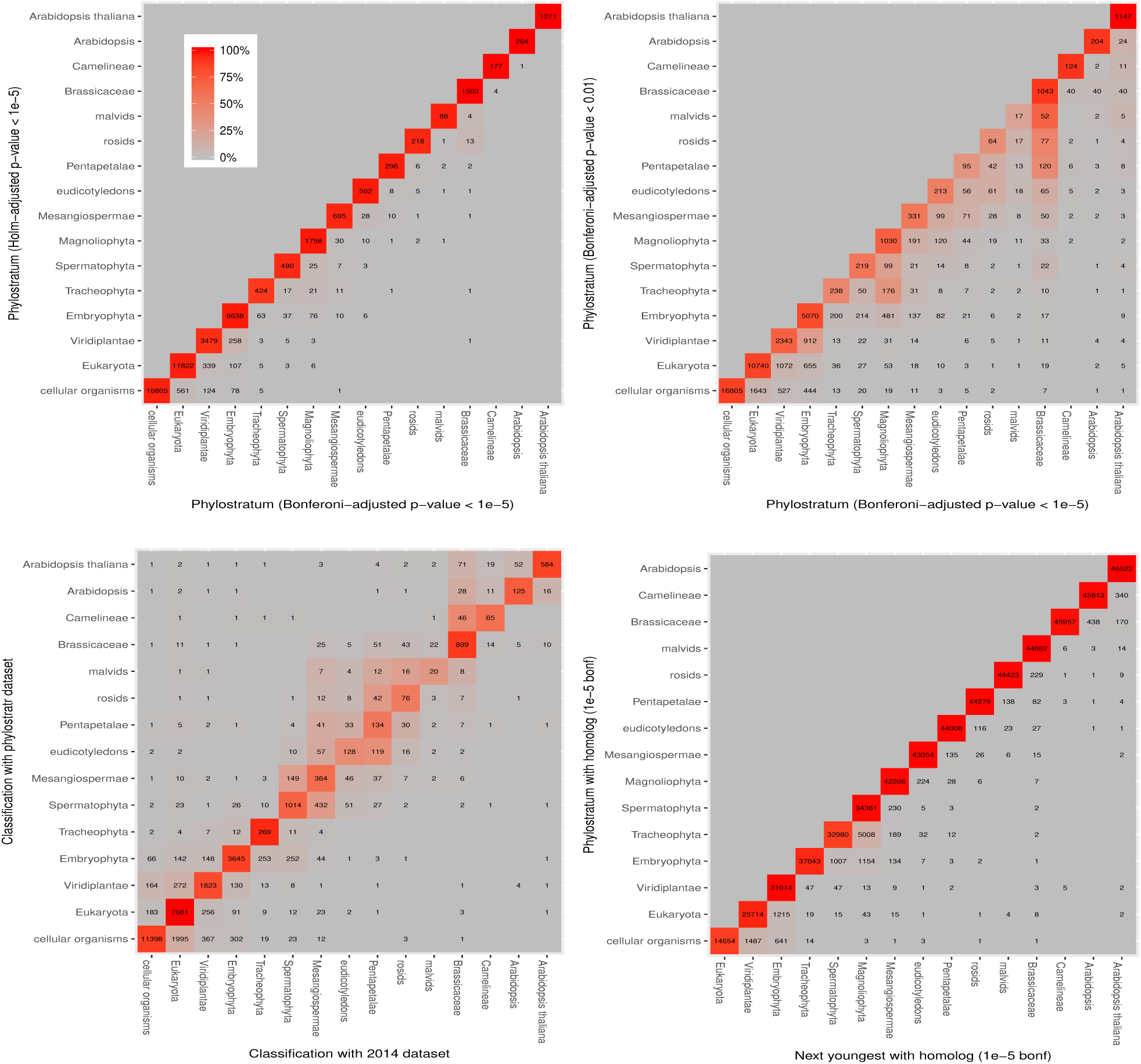
Heat map comparisons of phylostratigraphic analyses for *A. thaliana.* **(A)** Phylostratum classifications with P-values adjusted by Bonferroni versus Holm methods. The methods give near identical results; the Bonferroni method is more stringent, leading to a slight shift towards younger classifications. **(B)** Changing from a high Bonferroni cutoff (0.05) to a low cutoff (10^−5^) results in a shift towards younger classifications that is minor for some strata, but more significant for others; assignments to the *A. thaliana* stratum, the orphan genes, are very robust. **(C) phylostratr** results versus [5]. Classifications are similar between the two studies, the most notable difference is the several thousand more genes assigned to the cellular organisms stratum by `phylostratr`, likely because of a more diverse representation of prokaryotic species in the current study (85 vs 5 spp.). **(D)** The numbers of genes that have a significant (inferred) homolog in a phylostratum (Y axis) and their closest significant homolog in a younger phylostratum (X axis). For example, 1 of the 25,714 *A. thaliana* genes that genes that have a homolog in the Eukaryota stratum (Y axis) has no inferred homolog in any branch until rosids (X axis). Genes far from the diagonal are likely caused by false positives or potentially by lateral gene transfer.

Classifications into phylostrata are often quite robust against cutoff choice. For example, only 9.8% of the genes classified as orphan, *A. thaliana-specific*, under the 10^−5^ cutoff were classified into an older stratum when the cutoff was changed to 0.01. Other strata show larger changes, for example, 53 out of 78 of the genes tracing to the malvids stratum using a cutoff of 0.05 are assigned to the next-youngest stratum, Brassicaceae, if the cutoff lowered to (10^−5^) (see **Figure 2)**. This shows that most assignments to the malvids were based on fairly weak evidence.

This approach accounts for the differences in the number of species sampled from each phylostratum. This simplifies inter-study comparisons, and especially meta-analyses across evolutionary clades, which will become more important as we get better tools and more sequence data. A limitation of the approach, and one that is shared in all other BLAST-based phylostratigraphy analyses, is the assumption that the highest similarity scores for a focal species gene against the species of a phylostratum are independent. Considering dependencies is a problem grounded in gene evolution that is beyond the scope of this paper.

Homology could be inferred from data other than, or in addition to, the BLAST E-value. For example, the degree of sequence coverage (the percent of the query sequence that is covered by the alignment) could be incorporated into the homology decision. The user may provide a classifier function to customize this behavior.

### 3.7 Infer phylostrata from homology inferences

Given the location of inferred homologs across the phylogenetic tree, the phylostrata for each gene can be determined. In conventional phylostratigraphy, and in `phylostratr` currently, each gene is assigned to the most evolutionarily distant phylostratum that contains an inferred homolog. Alternatives approaches are explored in [24].

### 3.8 Build diagnostics and reports

`phylostratr` makes the parameters and results of the heavy computational steps easily accessible, providing a rich suite of tools for diagnostics and exploration (**Figure 1)**. Diagnostics include protein quality assessment (**Table S1**), organelle genome checking (**Table 2)**, heat maps of inference methods and of skipped clades **Figure 2,** and gene by clade tree-heat maps (**Figure 3)** are exemplified in the case studies presented in **Section 4**.

**Figure 3.**
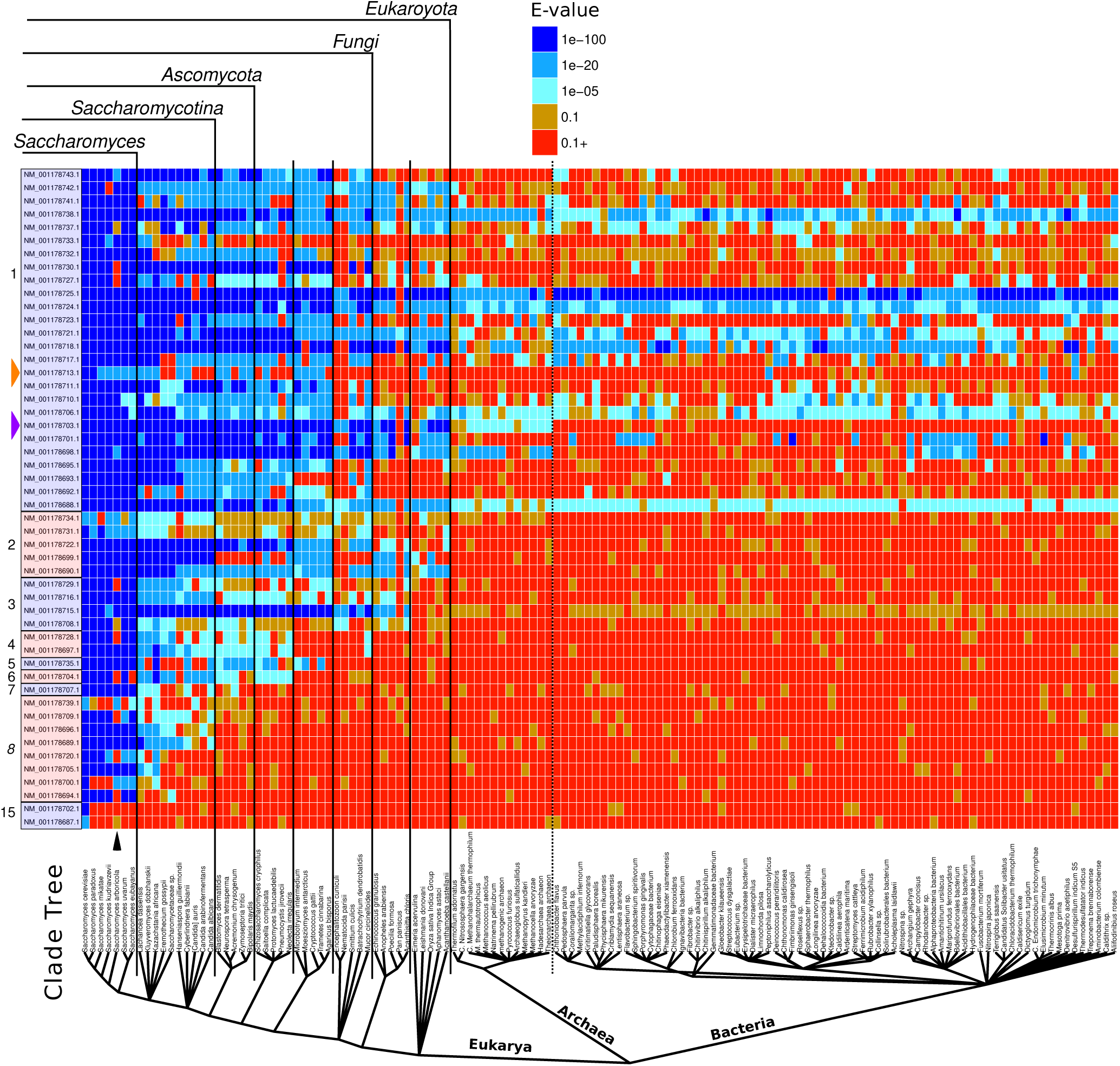
`phylostratr` gene by clade tree-heat map for *Saccharomyces cerevisiae.* The heat map shows the degree of similarly of each focal species gene (far left column) to that of its closest homolog in each species of the clade tree. Proteins encoded by consecutive *S. cereviciae* genes (NM_001178687.1 - NM_001178743.1), sorted by phylostratum membership, are compared to those of genomes of 132 species. The designated phylostratum membership (1-15) is indicated on the far left. Orange wedge: This gene has homologs in the fungal phylostrata, but not in other eukaryotes, and also has bacterial homologs; it is potentially associated with a horizontal transfer event. Magenta wedge: this gene is present in eukaryotes and archeae but not in bacteria. Black arrow: The nine “missing” homologs in *S. arboricola* could be due to an incomplete genome annotation or a biological phenomenon such as a rapidly evolving genome or massive gene loss; the statistics for the *S. arboricola* genome assembly at NCBI support the former explanation. This plot was built with `ggtree` [15] and annotated with Inkscape.

## 4 Results and Discussion

We used `phylostratr` for phylostratigraphic analyses of all annotated genes in two test cases: *Arabidopsis thaliana* and *Saccharomyces cerevisiae.* In both case studies, to illustrate diagnostic features of `phylostratr`, we intentionally retained a few species with low quality nuclear or organelle gene annotations. For *A. thaliana*, the diagnostics from an initial analysis enabled us to quickly realize that we needed a second iterations with expanded protein sequence data (see R code of **Figure 1)**. The full R scripts and diagnostic output for each case study are included in the Supplementary Material.

### 4.1 phylostratr comparison to published results

We compare the `phylostratr` results for *A. thaliana* to those from an earlier study in 2014 [5] (**Table 1)**. A more in depth comparison is shown in the `phylostratr` diagnostic of **Figure 2C**, which maps how gene assignments changed between the studies. This figure reveals that the relatively large differences between studies seen in the table are due mostly to transitions of assignments to closely-related strata.

**Table 1.**
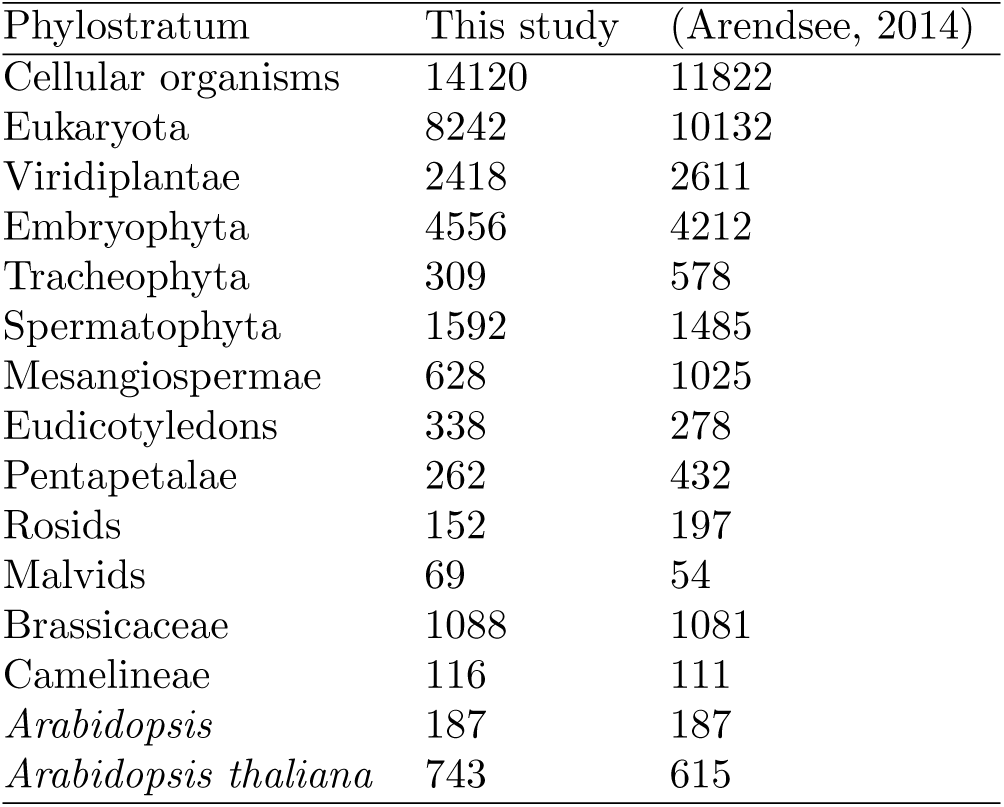
A comparison of the partitioning of genes between phylostrata in the current study and that of [5].

| Phylostratum | This study | (Arendsee, 2014) |
| --- | --- | --- |
| Cellular organisms | 14120 | 11822 |
| Eukaryota | 8242 | 10132 |
| Viridiplantae | 2418 | 2611 |
| Embryophyta | 4556 | 4212 |
| Tracheophyta | 309 | 578 |
| Spermatophyta | 1592 | 1485 |
| Mesangiospermae | 628 | 1025 |
| Eudicotyledons | 338 | 278 |
| Pentapetalae | 262 | 432 |
| Rosids | 152 | 197 |
| Malvids | 69 | 54 |
| Brassicaceae | 1088 | 1081 |
| Camelineae | 116 | 111 |
| *Arabidopsis* | 187 | 187 |
| *Arabidopsis thaliana* | 743 | 615 |

The major difference between the studies is the selection of species in the search space. The 2014 study used a manually-curated selection of representative species and a 10^−5^ E-value cutoff for hits against independent proteome databases. The current study uses automatically-selected UniProt proteomes, supplemented with a *Ginkgo* proteome, and a 10^−5^ nested, adjusted p-value cutoff. The 2014 study used the TAIR10 annotation of *A. thaliana*, whereas the current study uses Araport11 [25]. To account for the annotation differences, we used only the gene models common to both annotations. Also, we only compared the common strata. For example, the Brassicales branch was represented only in the 2014 study, so for the comparison, all its members were merged back into the more recent Brassicaceae branch.

In general, the phylostratigraphic classifications were similar in both studies. The differences are likely predominantly due to two factors: the expansion of the number of species included in the `phylostratr` study (85 prokaryotes versus 12) and, the increased quality and annotations of genomes in 2018 compared to 2013. Thus, several thousand more genes were traced back to an earlier origin by the `phylostratr` analysis.

There are three main advantages of using `phylostratr` for this analysis. First, the use of `phylostratr` removes the time-consuming manual step. Second, using `phylostratr` enables a researcher access to a range of detailed diagnostic plots, this can be valuable for the original researcher, or to a researcher who is comparing her results to the present study. Third, the `phylostratr` study is completely reproducible.

### 4.2 Phylogenetic tree, data collection

For the *A. thaliana* case study, we used the default NCBI common tree and UniProt proteomes for the target search space. We set a preference for reference proteomes (weight of 1.1) and designated a maximum of five representatives per phylostratum using the algorithm explained in **Section 3.1**. From the 1269 eukaryotic proteomes first retrieved in in UniProt, we filtered out a diverse set of 45 proteomes. We added human and *S. cerevesiae* genomes, because they are highly curated. After looking at the diagnostic plots from an initial run, we noted that the genome of the single representation of the Spermatophyta stratum in NCBI (*Picea glauca*) was sparsely annotated. To obviate this under-annotation, we added the transcriptome of the Spermatophyte *Ginkgo biloba* from the 1KP project [26–29] to the target search space.

The *S. cerevisiae* case study had the challenge that the genus node of the NCBI common tree for *Saccharomyces* was the immediate parent of around 100 species. To resolve differences between the species, we replace this branch with our own custom tree (**Figure S1**). The remainder of the study was similar to the *A. thaliana* case.

For both case studies, we selected a diverse set of 85 prokaryotic species, comprised of one species from each class of Bacteria and Archaea, sampled at random from the UniProt reference proteomes. Despite this sampling of prokaryotes, the diversity of the prokaryotes genomes is so great, their content so fluid, and their sequence so highly divergent from the eukaryotic focal species, that some genes with a prokaryotic origin will likely be missed. As noted in **Section 3.5,** using a more sensitive tool, such as HMMER, would be more suitable for very deep (eukaryote to prokaryote) similarity search.

### 4.3 Protein and organelle diagnostics

`phylostratr` provides functions that summarize and visualize proteome statistics. These can be used as another way to diagnose irregularities in proteome qualities. For example, in the *S. cerevisiae* study, the proteome of the primate *Pan paniscus* (a bonobo) is annotated as having only 802 genes (**Table S7**). The median protein length of 181 amino acids, is also far below that of the other species in the phylostratum. This suggests the genome is incomplete, poorly sequenced or poorly annotated and should not be included in the study.

For UniProt proteomes, `phylostratr` can determine which genes are encoded on organelle genomes. This information is particularly important in the phylostratigraphy of plant species, in which organelle genomes are large and, unfortunately, are often missing from the proteomes cataloged in UniProt. If the focal species contains organelle genomes, but the targets do not, the organelle genes can appear to be younger than they actually are. Further compounding this paucity of organellar proteomes, is the flux of genes from plastid to nuclear genomes. Genes that are transferred to the nuclear genome from the plastid genome, will appear to be young if chloroplast genomes are missing in the more ancient species that are part of the phylostratigraphic analysis. In the *A. thaliana* study, 19 of the 41 plant proteomes appear to be missing most or all mitochondrial sequences (see **Table 2** and **Table S2**).

**Table 2.**
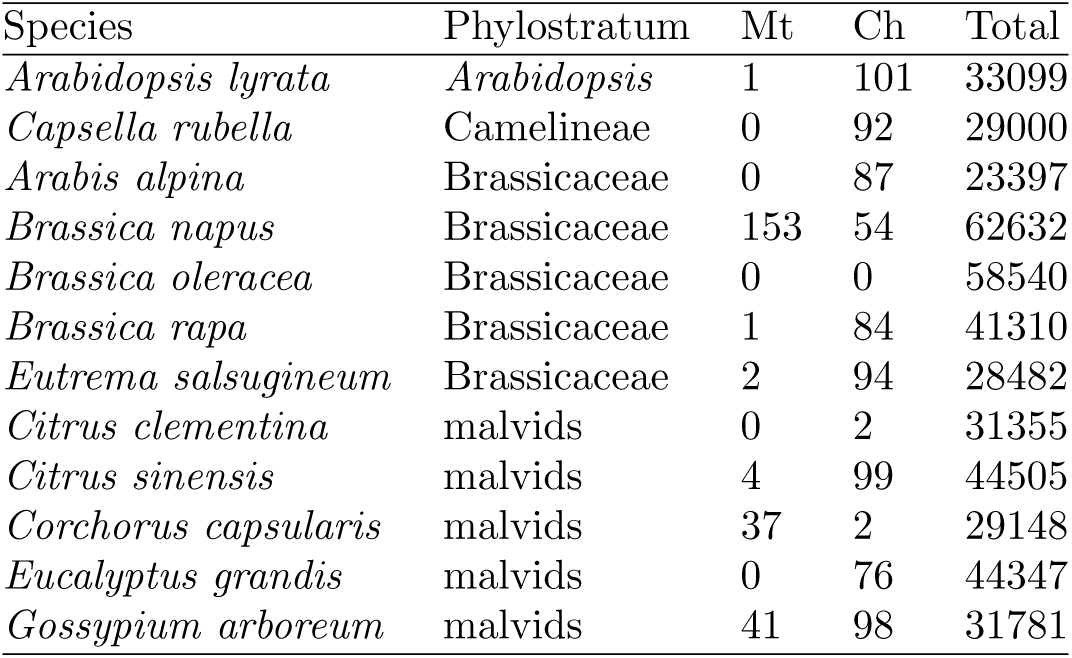
`phylostratr` diagnostics: mitochondrial, chloroplast and total UniProt **gene** counts for youngest four phylostrata in the *Arabidopsis thaliana* study. *A. thaliana* itself is not included, since a user-provided sequence, with no UniProt identiers, was used. See **Table S2** for the full list. **Mt**/**Ch** represent the number of **genes** represented in the given proteome that are annotated as transcribed from the mitochondrial or chloroplast genomes, respectively. Total represents all genes annotated in this species. **Total** is the total number of proteins included in the UniProt proteome.

### 4.4 Similarity, homology inference, and “gene by clade tree-heat map” diagnostics

The `phylostratr` default parameters were used for the BLAST and homology inference in both case studies. `phylostratr` provides graphical diagnostics from the similarity data (**Figure 3)**. Protein diagnostics for this species (discussed in **Section 4.3)**, which show *S. arboricola* has about a third fewer annotated proteins than the other species in the genus. Thus, in a subsequent analysis, we might want to remove this species and substitute another with a better annotated genome.

A gene by clade tree-heat map enables the user to quickly explore the scores of individual genes across target species. For the *S. cerevisiae* `phylostratr` analysis, one page of this diagnostic is show in **Figure 3)**. From these graphics, we can quickly see irregularities in the input data on a gene by gene basis. For example, the *Pan paniscus* genome shows a similarity to only one of the genes on this page of diagnostics: NM_001178721.1, which is a highly canonical serine/threonine-protein kinase, HAL4/SAT4.

The heatmaps can also be used to flag candidate genes for further investigation (**Figure 3B**). A researcher can quickly compare phylostratigraphic profiles across many genes to discover potential cases of horizontal transfer [30–32], gene loss, or other confounding histories. For example, a gene present only in a narrow eukaryotic clade, but with prokaryotic homologs, would be reported as a member of the phylostratum “cellular organisms”. However, the heatmap would reveal this gene as a candidate for horizontal transfer from a prokaryote prior to the speciation event leading to the eukaryotic clade in which it is found (an example is shown in **Figure 3)**.

### 4.5 Homology inference and quantitative diagnostics

Another approach to detect genes with irregular phylostratigraphic profiles is to find genes that “skip” strata (have no homologs in several intermediate strata, but do have homologs in more basal strata). These are summarized for *A. thaliana* (**Table S4**) and *S. cerevisiae* (**Table S9**). Technical or biological factors could explain this phenomenon. Genes that skip a stratum could result from poor quality data across all representatives of the stratum. For example, 10 of the 13 mitochondrial genes annotated in *S. cerevisiae* skip 4 or more strata; this is simply due to absence of mitochondrial genomes in many target genomes. Strata may also be skipped because of false positives, such as a non-specific match to a repetitive region. Alternatively, there could be a biological explanation, for example, multiple deletion events or a horizontal gene transfer.

**Figure 2D** is a diagnostic that visualizes the genes assigned to a given stratum and shows their next significant hit in the stratum. About 78 percent of the genes of the focal species *A. thaliana* have inferred homologs in a given strata, and then in each of the more closely-related strata. This can be seen from the genes indicated in the boxes on the diagonal; genes represented on the diagonal are those where no phylostratum is “skipped”. For example, 25204 *A. thaliana* genes have homologs within Eukaryota, and their next closest homolog is in the next youngest strata, Viridiplantae.

Genes represented in cells off the diagonal indicate cases where a deeper stratum has an inferred homolog, but adjacent shallower branches do not. About 25% of the genes in the *A. thaliana* study “skip” strata, i.e., have no inferred homolog in the next-youngest strata. For example, five genes have significant hits within Eukaryota and Brassicaceae strata, but have no inferred homologs in the intermediate strata. This could imply that either the ancient inferred homologs are false positives, the intermediate genomes are incomplete, or that the genes that skip phylostrata were laterally transferred to the ancestor of the Brassicaceae family. Lateral transfer is an especially interesting case, as seen in the 1487 genes assigned to cellular organisms that skip Eukaryota, and whose next significant hits are in Viridiplantae. These genes are predominantly associated with photosynthesis and presumably result from the major endosymbiotic event by which a Viridiplantae ancestor acquired chloroplasts.

## 5 Conclusions

The output of `phylostratr` can serve as the foundation for more specialized phylostratigraphic analyses. For example, the age estimates from `phylostratr` can be passed to the `myTAI` R package [33], and used to explore the transcriptomic “hourglass” developmental patterns. More generally, all the BLAST results, and all sequence data, can be trivially accessed for downstream analysis.

There are many avenues for future development of `phylostratr` by adding support for new algorithms such as: `HHblits` [34]; handling for estimation of error, as explored in [35]; and statistical analysis of evolutionary trends across time.

`phylostratr` is a highly automated and flexible platform for phylostratigraphy that bypasses tedious data handling and manipulation and provides an accessible, standardized framework for phylostratigraphic analysis. It provides a suite of tools that helps researchers avoid common pitfalls in phylostratigraphy, gain insight into the ontogenies of genes, and uncover the richness and complexity of gene evolution. Being an R package, `phylostratr` can be integrated naturally into R-based bioinformatics and statistical pipelines.

## Acknowledgements

We thank Manhoi Hur and Jennifer Chang for insightful discussions and encouragement. We also thank Scott Chamberlain and rOpenSci for help with `taxizedb`.

## References

[1] Domazet-Lošo, T. et al. (2007) A phylostratigraphy approach to uncover the genomic history of major adaptations in metazoan lineages. Trends in Genetics 23, 533–539

[2] Šestak, M.S. and Domazet-Lošo, T. (2014) Phylostratigraphic profiles in zebrafish uncover chordate origins of the vertebrate brain. Molecular biology and evolution 32, 299–312

[3] Bhandary, P. et al. (2017) Raising orphans from a metadata morass: a researcher’s guide to re-use of public’omics data. Plant Science

[4] Khalturin, K. et al. (2009) More than just orphans: are taxonomically-restricted genes important in evolution? Trends in Genetics 25, 404–413

[5] Arendsee, Z.W. et al. (2014) Coming of age: orphan genes in plants. Trends in plant science 19, 698–708

[6] Scott, G.K. et al. (1986) Fish antifreeze proteins: recent gene evolution. Canadian Journal of Fisheries and Aquatic Sciences 43, 1028–1034

[7] Ekstrom, A. and Yin, Y. (2016) Orfanfinder: automated identification of taxonomically restricted orphan genes. Bioinformatics 32, 2053–2055

[8] Cheng, X. et al. (2015) A “developmental hourglass” in fungi. Molecular biology and evolution 32, 1556–1566

[9] Consortium, U. et al. (2014) Uniprot: a hub for protein information. Nucleic acids research p. gku989

[10] Team, R.C. (2000) R language definition. Vienna, Austria: R foundation for statistical computing

[11] Johnson, M. et al. (2008) Ncbi blast: a better web interface. Nucleic acids research 36, W5–W9

[12] Chamberlain, S. et al. (2018). ropensci/taxizedb: taxizedb v0.1.6

[13] Wickham, H. et al. (2017) dplyr: A Grammar of Data Manipulation. R package version 0.7.4

[14] Wickham, H. (2009) ggplot2: Elegant Graphics for Data Analysis. Springer-Verlag New York

[15] Yu, G. et al. (2017) ggtree: an r package for visualization and annotation of phylogenetic trees with their covariates and other associated data. Methods in Ecology and Evolution 8, 28–36

[16] Federhen, S. (2011) The ncbi taxonomy database. Nucleic acids research 40, D136–D143

[17] Finn, R.D. et al. (2015) Hmmer web server: 2015 update. Nucleic Acids Research 43, W30–W38

[18] Domazet-Lošo, T. et al. (2017) No evidence for phylostratigraphic bias impacting inferences on patterns of gene emergence and evolution. Molecular biology and evolution 34, 843–856

[19] Moyers, B.A. and Zhang, J. (2014) Phylostratigraphic bias creates spurious patterns of genome evolution. Molecular biology and evolution 32, 258–267

[20] Moyers, B.A. and Zhang, J. (2016) Evaluating phylostratigraphic evidence for widespread de novo gene birth in genome evolution. Molecular biology and evolution 33, 1245–1256

[21] Moyers, B.A. and Zhang, J. (2017) Further simulations and analyses demonstrate open problems of phylostratigraphy. Genome Biology and Evolution

[22] Stancik, I.A. et al. (2018) Serine/threonine protein kinases from bacteria, archaea and eukarya share a common evolutionary origin deeply rooted in the tree of life. Journal of molecular biology 430, 27–32

[23] Moyers, B. (2017) On gene age, gene origins, and evolutionary trends

[24] Capra, J.A. et al. (2012) Proteinhistorian: tools for the comparative analysis of eukaryote protein origin. PLoS computational biology 8, e1002567

[25] Cheng, C.Y. et al. (2017) Araport11: a complete reannotation of the arabidopsis thaliana reference genome. The Plant Journal 89, 789–804

[26] Wickett, N.J. et al. (2014) Phylotranscriptomic analysis of the origin and early diversification of land plants. Proceedings of the National Academy of Sciences 111, E4859–E4868

[27] Matasci, N. et al. (2014) Data access for the 1,000 plants (1kp) project. Gigascience 3, 17

[28] Xie, Y. et al. (2014) Soapdenovo-trans: de novo transcriptome assembly with short rna-seq reads. Bioinformatics 30, 1660–1666

[29] Johnson, M.T. et al. (2012) Evaluating methods for isolating total rna and predicting the success of sequencing phylogenetically diverse plant transcriptomes. PloS one 7, e50226

[30] Méheust, R. et al. (2016) Protein networks identify novel symbiogenetic genes resulting from plastid endosymbiosis. Proceedings of the National Academy of Sciences 113, 3579–3584

[31] Yue, J. et al. (2012) Widespread impact of horizontal gene transfer on plant colonization of land. Nature communications 3, 1152

[32] Bock, R. (2017) Witnessing genome evolution: Experimental reconstruction of endosymbiotic and horizontal gene transfer. Annual review of genetics 51

[33] Drost, H.G. et al. (2017) mytai: Evolutionary transcriptomics with r. Bioinformatics

[34] Remmert, M. et al. (2012) Hhblits: lightning-fast iterative protein sequence searching by hmm-hmm alignment. Nature methods 9, 173–175

[35] Liebeskind, B.J. et al. (2016) Towards consensus gene ages. Genome biology and evolution 8, 1812–1823

